# Identifying targetable metabolic dependencies across colorectal cancer progression

**DOI:** 10.1101/2022.03.23.485483

**Authors:** Danny N. Legge, Ewelina Stanko, Amy K. Holt, Caroline J. Bull, Tracey J. Collard, Madhu Kollareddy, Jake Bellamy, Sarah Groves, Eric H. Ma, Emma Hazelwood, David Qualtrough, Borko Amulic, Karim Malik, Ann C. Williams, Nicholas Jones, Emma E. Vincent

**Affiliations:** School of Translational Health Sciences, Dorothy Hodgkin Building, University of Bristol, Bristol, BS1 3NY, UK; School of Cellular & Molecular Medicine, University of Bristol; Integrative Epidemiology Unit, School of Population Health Science, University of Bristol; School of Biochemistry, University of Bristol; Metabolic and Nutritional Programming, Center for Cancer and Cell Biology, Van Andel Institute; Faculty of Health and Life Sciences, University of the West of England; Institute of Life Science, Swansea University Medical School, Swansea University, SA2 8PP

**Author notes:** Corresponding author Emma E. Vincent, Dorothy Hodgkin Building, Whitson Street, Bristol, UK, BS1 3NY.

## Abstract

Colorectal cancer (CRC) is a multi-stage process initiated through the formation of a benign adenoma, progressing to an invasive carcinoma and finally metastatic spread. Tumour cells must adapt their metabolism to support the energetic and biosynthetic demands associated with disease progression. As such, targeting cancer cell metabolism is a promising therapeutic avenue in CRC. However, to identify tractable nodes of metabolic vulnerability specific to CRC stage, we must understand how metabolism changes during CRC development. Here, we use a unique model system – comprising human early adenoma to late adenocarcinoma. We show that adenoma cells transition to elevated glycolysis at the early stages of tumour progression but maintain oxidative metabolism. Progressed adenocarcinoma cells rely more on glutamine-derived carbon to fuel the TCA cycle, whereas glycolysis and TCA cycle activity remain tightly coupled in early adenoma cells. Adenocarcinoma cells are more flexible with respect to fuel source, enabling them to proliferate in nutrient-poor environments. Despite this plasticity, we identify asparagine (ASN) synthesis as a node of metabolic vulnerability in late-stage adenocarcinoma cells. We show that loss of asparagine synthetase (ASNS) blocks their proliferation, whereas early adenoma cells are largely resistant to ASN deprivation. Mechanistically, we show that late-stage adenocarcinoma cells are dependent on ASNS to support mTORC1 signalling and maximal glycolytic and oxidative capacity. Resistance to ASNS loss in early adenoma cells is likely due to a feedback loop, absent in late-stage cells, allowing them to sense and regulate ASN levels and supplement ASN by autophagy. Together, our study defines metabolic changes during CRC development and highlights ASN synthesis as a targetable metabolic vulnerability in later stage disease.

## Introduction

Colorectal cancer (CRC) ranks second globally with regards to cancer mortality, due to limited treatment options and low response rates (1, 2). This, coupled with the concerning rise of CRC in younger people (3), highlights an urgent need for further understanding of the underlying molecular mechanisms to improve therapeutic options. Despite the molecular genetic progression of the adenoma-carcinoma sequence being well defined (4, 5), the underlying metabolic changes that accompany this progression have not been well characterised. As targeting tumour cell metabolism is a promising therapeutic avenue, defining metabolic changes across tumour progression has the potential to highlight novel vulnerabilities of CRC cells.

A number of recent *in vivo* studies across different tumour sites have revealed that cancer cells transition through diverse metabolic states at different stages of development, presenting potential opportunities to target these distinct metabolic vulnerabilities (6). However, metabolic plasticity is a major clinical barrier, with drugs targeting this plasticity the subject of ongoing clinical trials (7). Identification of metabolic dependencies at specific stages of progression is crucial for designing more effective treatments. Combination therapies appear to be most effective when targeting metabolic vulnerabilities. For instance, a recent study designed to identify strategies to target metabolic plasticity found that upon deletion of glutaminase (GLS) 1 and 2, liver tumours rewire central carbon metabolism via compensatory action of transamidases, requiring targeting of both pathways to inhibit tumour cell proliferation (8).

To target specific stages of CRC, including the pre-malignant or very earliest stages of cancer development, we must first understand how these stages are characterised with respect to metabolism, which requires an appropriate model of colorectal origin. *In vitro* malignant progression models of cancer have potential to be powerful tools for identifying metabolic changes across tumour progression, but availability of these remain limited. These models have the advantage of being of human origin and enable the investigation of cellular metabolic activity using techniques such as stable isotope labelling (SIL). A recent study used transformed normal human lung fibroblasts to produce a progression model of pre-malignant to highly tumorigenic cell lines (9). Large-scale proteomic and metabolomic analyses revealed perturbations in nitrogen metabolism across the series, implicating a high phosphoribosyl pyrophosphate amidotransferase (PPAT)/GLS1 ratio in small cell lung cancer and other malignancies through increased nucleotide synthesis (9). Another study characterised a metabolic signature of breast cancer progression, demonstrating reduced oxidative metabolism and glycolysis maintenance across cell line progression (10). These findings highlight the potential of *in vitro* malignant progression models and emphasise the need for cancer-specific models to understand the nuances of metabolic reprogramming in tumours arising from different tissues.

Despite the lack of *in vitro* models of CRC development, attempts have been made to investigate the metabolic profile of CRC tissue at the different stages of disease progression for use in prognostics, diagnostics and therapeutics (11, 12). One such study, using clinical samples from four cohorts across two countries, discovered a signature of 15 metabolites which could be used to predict recurrence and survival, despite the variable mutations, genetic backgrounds and disease stage (11). Identification of metabolic features specific to the pre-malignant stage may be key for CRC prevention. Increased consumption of glucose and inositol have been reported in polyps (13); however, exploration of dynamic metabolic activity in adenoma cells, rather than measurement of static metabolite levels alone, is largely lacking.

Here, we identify metabolic changes across CRC development to reveal tractable nodes of metabolic vulnerability with future potential for CRC treatment. To do this we use a unique model system derived from a human colorectal adenoma, transformed in sequence and recapitulating the genetic changes seen *in vivo*, allowing us to probe the different stages of colorectal tumour progression. We show that the metabolism of early adenoma cells differs considerably from that of adenocarcinoma cells, revealing metabolic changes across CRC development. Early adenoma cells favour oxidative metabolism and are not as metabolically plastic; but are less dependent on *de novo* asparagine (ASN) biosynthesis than their progressed counterparts. Using our approach, we discovered that the ASN synthesis pathway plays an important role in late-stage adenocarcinoma cells. This is supported by clinical data revealing *ASNS* expression is elevated in colorectal tumour and metastatic tissue in comparison to normal tissue and is associated with poorer overall survival. Loss of ASNS blocks proliferation in late-stage cells, attenuating mTORC1 signalling and ATP production, whereas early adenoma cells are largely resistant to *ASNS* knockdown due to their ability to sense ASN levels and source it through autophagy. Thus, using our progression model of CRC, we reveal that ASNS is a node of metabolic vulnerability in CRC cells. Our results highlight ASNS as a promising therapeutic target to combat aggressive tumour cells while shielding surrounding healthy tissue. Hence, ASNS dependence represents a weakness in the metabolic programme of late CRC and may be an attractive future therapeutic target.

## Results

### Glycolytic and oxidative metabolism shift early in colorectal tumour progression

To investigate metabolic changes across colorectal tumour progression, we have used a model system consisting of four cell lines of the same lineage (Figure 1A). Serial transformation of this cell line series has been described in detail previously (14). Briefly, each cell line represents a different stage in colorectal cancer progression and has been demonstrated to model human colorectal carcinogenesis *in vivo* (14). The original cell line (PC/AA) was derived from a human colorectal adenoma and cultured *in vitro* to produce the clonogenic, anchorage-dependent line; AA/C1 (referred to henceforth as C1). The AA/C1/SB (SB) cell line was generated from C1 cells following sodium butyrate treatment. These cells exhibited anchorage independent growth, a higher proliferative rate and increased colony forming efficiency (CFE) but were unable to initiate tumours in nude mice. The SB cells were subsequently treated with the carcinogen N-methyl-N’- nitro-N-nitrosoguanidine (15) to generate the AA/C1/SB10 (10C) adenocarcinoma line which exhibited further increased CFE, anchorage independence and tumour initiation when xenografted to nude mice. Finally, cells were extracted from 10C xenograft tumours and further cultured *in vitro*, giving rise to the AA/C1/SB/10C/M (M) cell line. M cells showed increased proliferation, CFE and tumour initiating capacity in xenograft experiments compared to 10C cells and therefore represent a highly tumorigenic, late-stage adenocarcinoma (14).

**Figure 1.**
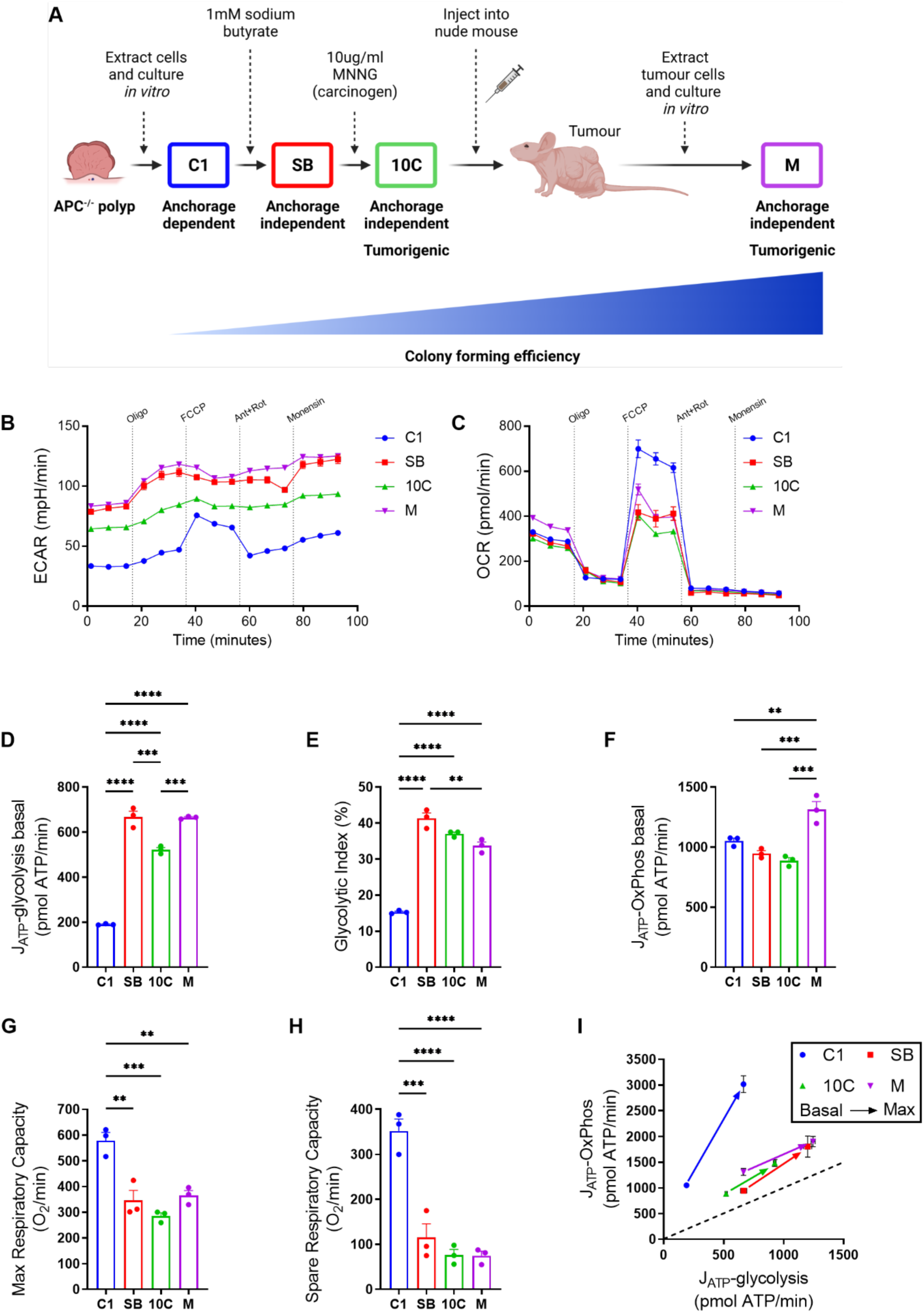
**A bioenergetic shift occurs early in colorectal tumour progression** (A) Schematic showing generation of the *in vitro* colorectal adenoma to carcinoma progression model. Figure created using BioRender.com. (B & C) Seahorse Extracellular Flux Analyzer traces of extracellular acidification rate (ECAR; B) and oxygen consumption rate (OCR; C). Oligo, oligomycin A; FCCP, carbonyl cyanide p-trifluoro-methoxyphenyl hydrazone; Ant, antimycin A; Rot, rotenone. (D) Rate of ATP generation via glycolysis at baseline. (E) Glycolytic index; ATP produced by glycolysis as a percentage of the total ATP produced by the cell at baseline. (F) Rate of ATP generation via oxidative phosphorylation (OxPhos) at baseline. (G) Maximal respiratory capacity (MRC); OCR following FCCP addition. (H) Spare respiratory capacity (SRC); difference between baseline OCR and MRC. (I) Bioenergetic scope of tumour cell lines detailing shift from basal to maximal ATP production from glycolytic (J_ATP_-glycolysis) and oxidative (J_ATP_-OxPhos) metabolism. (B-I) Data are represented as mean ± SEM of three independent cultures. (D-H) Tukey’s multiple comparisons test. \*\**p*<0.01; \*\*\**p*<0.001; \*\*\*\**p*<0.0001.

Initially, to interrogate any overall differences in oxidative or glycolytic metabolism across progression, we performed a mitochondrial stress test using a Seahorse Extracellular Flux Analyzer to measure extracellular acidification rate (ECAR) and oxygen consumption rate (OCR) in each of the cell lines (Figure 1B and C).

Glycolytic rate was increased substantially in SB cells in comparison to C1, with no further increase in the 10C and M cell lines (Figure 1D), suggesting the switch to an increased glycolytic rate occurs early in colorectal tumour progression. Consistent with this, the C1 early adenoma cells have a significantly lower glycolytic index compared to the rest of the series (Figure 1E), representing the ATP produced by glycolysis as a percentage of the total ATP produced by the cell (16, 17).

Regarding oxidative phosphorylation (OxPhos), the most progressed adenocarcinoma cells (M) generate significantly more ATP through this pathway than the earlier stage cells, whereas there are no significant differences in ATP production between the C1, SB and 10C cells at baseline (Figure 1F). However, the early adenoma C1 cells have a much greater maximal (Figure 1G) and spare respiratory capacity (Figure 1H), suggesting these cells may have greater capability to increase ATP production through OxPhos when stressed. This could be a pro-survival adaptation of adenoma cells that comes at a fitness cost, subsequently lost during tumour progression. The shift from basal to maximal ATP production via glycolytic and oxidative metabolism in each of the cell lines in the progression reveals a distinct clustering of the SB, 10C and M cell lines, relative to the early stage C1 adenoma cells (Figure 1I).

To determine if the differences in oxidative metabolism are due to changes in mitochondrial number or morphology, we carried out transmission electron microscopy (TEM) to measure mitochondrial number, area, length, and roundness in our cell line series (Supplementary Figure S1A-G). These analyses revealed no significant differences in any of the parameters measured throughout the adenoma to carcinoma progression. We also assessed expression levels of the individual respiratory complexes by immunoblot (Supplementary Figure S1H). Although we observe some differences across the series, the most notable being increased expression of complex III in SB, 10C and M in comparison to C1, they are unlikely to explain the measured changes in OxPhos that we observe.

Taken together, these data indicate that increased glycolytic rate is an early event in colorectal tumour progression; however, this does not come at the expense of OxPhos, which is maintained and eventually increased in late-stage cells. These observations are in line with previous findings from *in vivo* analyses of human tumours originating from multiple tissue types (18), supporting the validity of our model.

### Contributions of glucose and glutamine to central carbon metabolism changes across colorectal adenoma to carcinoma progression

Central carbon metabolism is aberrant in cancer cells compared to normal tissue, with increased use of both glucose and glutamine by tumours being well-documented (19, 20). To explore in detail how central carbon metabolism is altered across colorectal cancer progression, we cultured our cell line series in the presence of uniformly labelled ^13^C-glucose (U-[^13^C]-Glc) or ^13^C-glutamine (U-[^13^C]-Q). Conventional metabolism of U- [13C]-Glc and U-[^13^C]-Q in tumour cells is illustrated in Supplementary Figure S2A and B. In general, we observed a decrease in incorporation of glucose-derived carbon into TCA cycle intermediates and associated non-essential amino acids (NEAA) in the more progressed lines in comparison to the C1 early adenoma cells, with the exception of α-ketoglutarate, fumarate and asparagine (Figure 2A). In contrast, we observed an increase in incorporation of glutamine-derived carbon into the TCA cycle and associated NEAAs in the more progressed tumour cells compared to early adenoma cells (Figure 2B). These findings are supported by mass isotopologue distribution (MID) analysis which confirm decreased abundance of fully labelled isotopologues from glucose (Figure 2C) and an increased abundance of the m+4 isotopologue from glutamine in TCA cycle metabolites (Figure 2D) in the more progressed adenoma and adenocarcinoma cells in comparison to the C1 early adenoma cells. A higher ratio of m+5 glutamate/m+5 glutamine in the more progressed lines in comparison to the early adenoma C1 cells from U-[^13^C]-Q also indicates reduced rates of glutaminolysis in C1 cells (Figure 2E).

**Figure 2.**
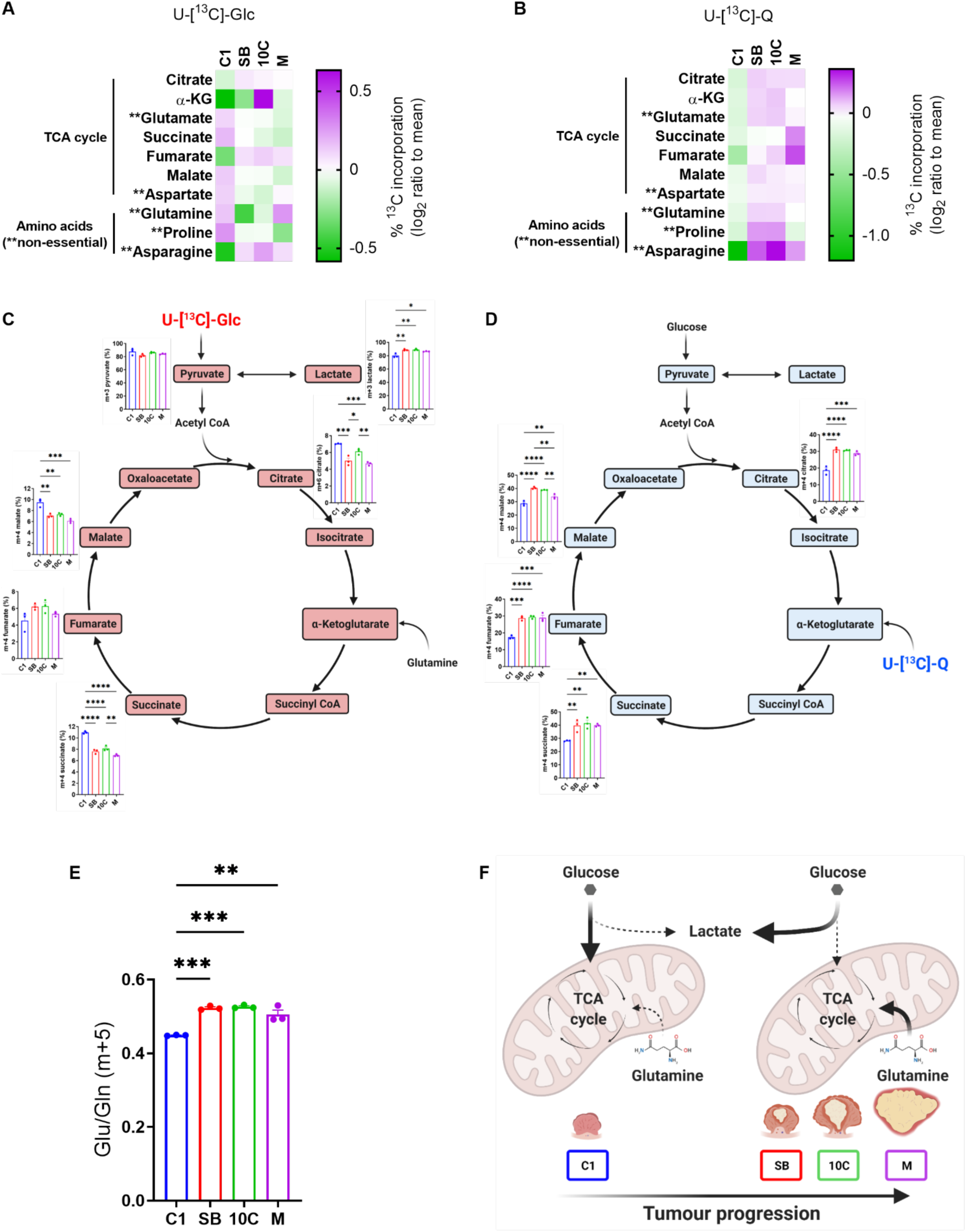
**Central carbon metabolism changes across colorectal tumour progression** (A & B) Heatmaps showing percentage incorporation of uniformly labelled U-[^13^C]-Glc (A) or U-[^13^C]-Q (B) into downstream metabolites in each cell line. Data are represented as log_2_ ratio to mean for each individual metabolite from three independent cultures. (C & D) Mass isotopologue distribution (MID) analysis of TCA cycle intermediates from U- [^13^C]-Glc (C) or U-[^13^C]-Q (D). (E) Ratio of m+5 glutamate/m+5 glutamine taken from U-[^13^C]-Q labelling data generated in (B & D). (F) Schematic outlining early shift in central carbon metabolism across colorectal adenoma to carcinoma progression. There is an early shift to enhanced glycolytic metabolism (lactate production; Figure 1) concomitant with increased glutamine-fuelled TCA cycle anaplerosis (Figure 2) as early adenoma (C1) cells transition to intermediate adenoma (SB) cells. This is maintained through the later stages of tumour progression to late adenocarcinoma. (C, D & F) Figures created using BioRender.com. (C-E) Data are represented as mean ± SEM of three independent cultures. Tukey’s multiple comparisons test. \**p*<0.05; \*\**p*<0.01; \*\*\**p*<0.001; \*\*\*\**p*<0.0001.

These observations, taken together with the bioenergetic analysis (Figure 1), suggest that in early adenoma C1 cells, glycolysis and the TCA cycle are more tightly coupled; exemplified by less lactate production (i.e., lower ECAR) through the glycolytic pathway (Figure 1) and more glucose carbon entering the TCA cycle (Figure 2). However, as cells progress from the early adenoma stage, the contribution of glucose carbon to the TCA cycle drops and lactate production is increased (i.e., higher ECAR; Figure 1). This is combined with increased use of glutamine-derived carbon for TCA cycle anaplerosis and non-essential amino acid (NEAA) biosynthesis (Figure 2). The metabolic switch happens early in the adenoma to carcinoma progression sequence, as SB cells are metabolically distinct from the C1 cells (summarised in Figure 2F).

### Late-stage colorectal adenocarcinoma cells acquire metabolic plasticity and resistance to nutrient stress

As a tumour grows and the cells acquire mutations and progress, the metabolic niche they inhabit becomes more challenging and competitive (21). Our study so far has identified changes in bioenergetics and central carbon metabolism across CRC development, we next wanted to understand if these changes support tumour progression during periods of nutrient stress.

To challenge the cells and test their metabolic flexibility, we carried out proliferation assays and extracellular flux analysis in low nutrient conditions (4 mM glucose or 0.5 mM glutamine). Relatively, the most progressed cells (M) proliferated most efficiently across 7 days in 4 mM glucose (Figure 3A-C). Interestingly, the SB and 10C cells performed worst in this assay, this could be explained due to the significant shift to glycolytic metabolism we observed in these cells in comparison to the early C1 cells (Figure 1D and E), making them more dependent on glucose availability.

**Figure 3.**
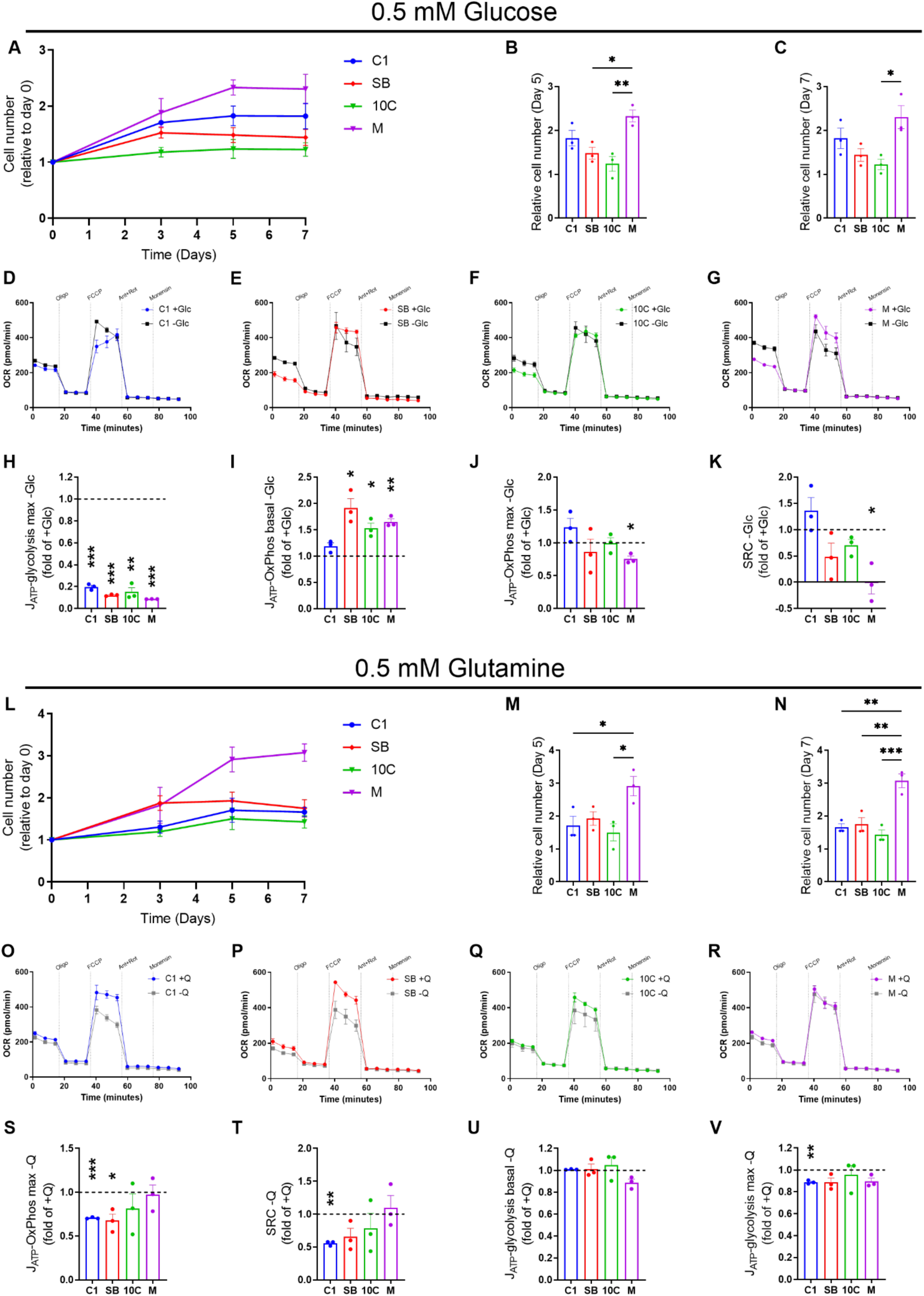
**Adaptation to nutrient stress varies across colorectal tumour progression** (A-C) Proliferation assay in low glucose conditions. (D-G) Seahorse Extracellular Flux Analyzer oxygen consumption rate (OCR) traces in C1 (D), SB (E), 10C (F) and M (G) cells in presence (+Glc; 10 mM) or absence (-Glc; 0 mM) of glucose. Oligo, oligomycin A; FCCP, carbonyl cyanide p-trifluoro-methoxyphenyl hydrazone; Ant, antimycin A; Rot, rotenone. (H-J) Rates of maximal ATP generation via glycolysis (H), basal ATP generation via OxPhos (I) and maximal ATP generation via OxPhos (J) in the absence of glucose. (K) Spare respiratory capacity (SRC) in the absence of glucose. Data expressed as -Glc/+Glc. (L-N) Proliferation assay low glutamine conditions. (O-R) OCR traces in C1 (O), SB (P), 10C (Q) and M (R) cells in presence (+Q; 2 mM) or absence (-Q; 0 mM) of glutamine. (S-V) Rate of maximal ATP generation via OxPhos (S), spare respiratory capacity (T), and rates of basal ATP generation via glycolysis (U) and maximal ATP generation via glycolysis (V) in the absence of glutamine. Data expressed as -Q/+Q. Data are represented as mean ± SEM of three independent cultures. (B, C, M & N) Tukey’s multiple comparisons test. (H-K; S-V) One sample t-test. \**p*<0.05; \*\**p*<0.01; \*\*\**p*<0.001.

To understand how colorectal tumour cells adapt bioenergetically to low glucose conditions, we performed a mitochondrial stress test in the absence of glucose in our tumour progression model (Figure 3D-G and Supplementary Figure S3A-D). Inhibition of glucose metabolism was confirmed by ∼80% decrease in maximal ATP production from glycolysis in all cell lines (Figure 3H and Supplementary Figure S3A-D). Interestingly, the C1 early adenoma cells were unable to increase baseline ATP production via OxPhos to compensate for reduced glycolytic ATP production following glucose restriction, in contrast to the more progressed cells (Figure 3I). As discussed previously, this may be explained by the relatively low levels of ATP production from glycolysis in the C1 cells, meaning less compensatory ATP production via OxPhos is required to maintain their bioenergetic state compared to the more progressed cells. Despite their ability to increase OxPhos at baseline in response to glucose restriction, the advanced adenocarcinoma cells (M) showed significantly reduced maximal ATP production via OxPhos (Figure 3J) and spare respiratory capacity (SRC) following FCCP addition in the absence of glucose (Figure 3K). This is an intriguing observation, though it does not appear to negatively impact their ability to survive and proliferate in low glucose conditions (Figure 3A-C).

Across 7 days in 0.5 mM glutamine, the most advanced adenocarcinoma cells (M) were again found to proliferate most efficiently, whereas there were no significant differences between the C1, SB and 10C cells (Figure 3L-N). We repeated the mitochondrial stress test assay in the absence of glutamine, these analyses revealed no change in basal OCR under glutamine restriction in any of the tumour cell lines (Figure 3O-R).

However, we found that the adenoma cells (C1 and SB) exhibited decreased levels of maximum OCR following FCCP addition in the absence of glutamine, which translated to significantly reduced ATP production (Figure 3S). This also resulted in decreased SRC in C1 cells (Figure 3T). Whilst there were no changes in basal ECAR in any of the tumour cells following glutamine restriction (Figure 3U), the maximal glycolytic rate was altered only in the C1 cells, where it was significantly reduced (Figure 3V and Supplementary Figure S3E-H). These data indicate that the early adenoma cells are no longer able to maintain maximal ATP production through OxPhos during glutamine restriction in the same way as the more progressed tumour cells.

Taken together, our analyses under nutrient-depleted conditions reveal that late-stage adenocarcinoma cells are more metabolically plastic and better able to adapt to low nutrient conditions than earlier stage adenoma and adenocarcinoma cells, translating to an increased ability to proliferate in low glucose and glutamine environments.

### Amino acid metabolism is dysregulated across colorectal adenoma to carcinoma progression

Thus far, our work has characterised significant metabolic reprogramming in the progressed adenocarcinoma cells in comparison to the early adenoma cell line (C1). To further understand these metabolic changes, we carried out proteomic analyses in the C1 and M cell line models; these lines representing the extremes of our colorectal tumour progression model. We identified significantly regulated expression of 2251 proteins between the cell lines (p<0.05; fold change>1.4; FDR<5%; Figure 4A). We performed over-representation analyses of these data using gene ontology (GO) biological process categorisation, which revealed ‘metabolic process’ as the top hit (Figure 4B). Further interrogation of these data using the KEGG pathway database established multiple metabolism-associated hits in the most highly enriched pathways. Hits included ‘valine, leucine and isoleucine degradation’, ‘propanoate metabolism’, ‘biosynthesis of amino acids’, ‘carbon metabolism’ and ‘metabolic pathways’ (Figure 4C). These findings highlight potential differences in amino acid metabolism between the early adenoma and late adenocarcinoma cells.

**Figure 4.**
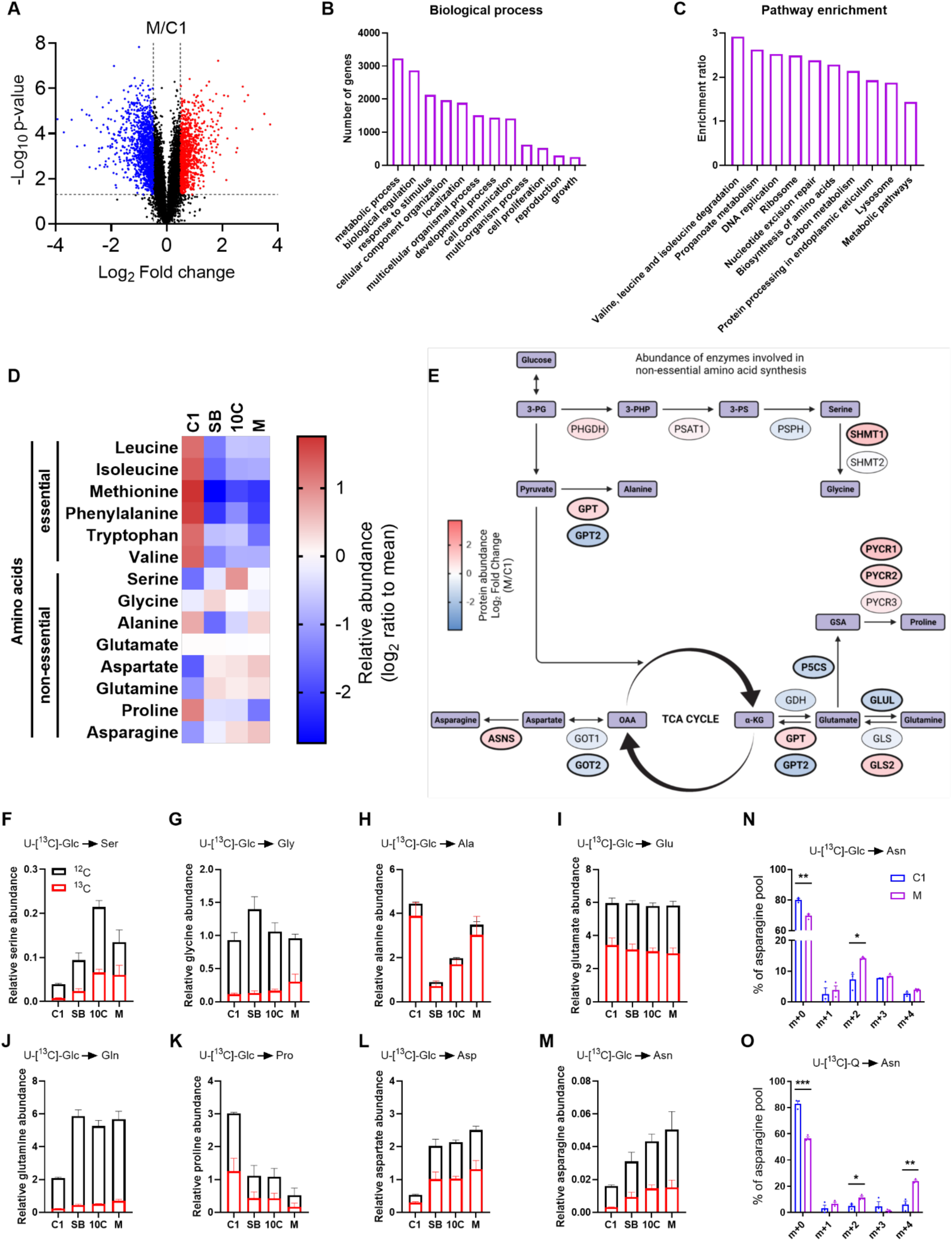
**Amino acid metabolism is dysregulated across colorectal adenoma to carcinoma progression** (A) Volcano plot of data summarising proteomic analysis of C1 vs M cells. Significantly upregulated proteins (M/C1) are marked in red, downregulated in blue and non-significant in black (FDR<5%, -log_10_ p-value>1.3 and log_2_ fold change+/- 0.48). (B & C) Gene Ontology (GO) biological process (B) and Kyoto Encyclopedia of Genes and Genomes (KEGG) pathway enrichment (C) analyses of the 2251 significantly regulated proteins identified in (A). (D) Heatmap displaying relative abundance of amino acids in each of the cell lines. Data are represented as log_2_ ratio to mean for each individual amino acid from three independent cultures. (E) Schematic of non-essential amino acid (NEAA) biosynthesis pathways branching off from glycolysis and the TCA cycle. Colour of enzyme (oval) represents degree of upregulation (red) or downregulation (blue) of abundance (M/C1) using proteomics data generated in (A). Significantly regulated enzymes in bold. Figure created using BioRender.com. (F-M) Relative abundance data of indicated NEAA in (D) including U-[^13^C]- Glc incorporation. (N & O) Mass isotopologue distribution of asparagine in C1 and M cells by U-[^13^C]-Glc (N) or U-^13^C- Q (O) labelling. Student’s t-test. \**p*<0.05; \*\**p*<0.01; \*\*\**p*<0.001. (F-O) Data are represented as mean ± SEM of three independent cultures.

Therefore, we next measured the abundances of amino acids across colorectal tumour progression in all four cell lines. We observed higher levels of all essential amino acids in C1 cells compared to the more progressed cancer cells (Figure 4D), likely due to increased usage in the more progressed tumour cells. This is consistent with the most highly enriched KEGG pathway in M cells - ‘valine, leucine and isoleucine degradation’ – in comparison with C1 cells (Figure 4C). In general, levels of NEAA are elevated in the more progressed cell lines in comparison to the C1 cells, with the exception of alanine and proline (Figure 4D). To explore these differences in more detail, we assessed abundance of enzymes involved in NEAA synthesis, taken from our earlier proteomics analysis (Figure 4A). Several enzymes involved in NEAA biosynthesis were significantly regulated (highlighted in bold) in late-stage adenocarcinoma cells (M) in comparison to early adenoma cells (C1), such as upregulation of GPT and GLS2, and downregulation of GOT2 and GPT2 (Figure 4E).

To investigate NEAA biosynthesis across colorectal tumour progression further, we used SIL in all four cell lines. Generally, abundance of NEAA and ^13^C incorporation (U-[^13^C]-Glc or U-[^13^C]-Q depending on the pathway being examined) was increased in the more progressed cells relative to the C1 early adenoma cells (Figures 4F-M), with the notable exceptions of alanine (Figure 4H) and proline (Figure 4K). Interestingly, we observed a stepwise increase in both aspartate (Figure 4L) and asparagine (Figure 4M) abundance (and U- [^13^C]-Glc incorporation) with progression from early adenoma to late adenocarcinoma. MID analysis confirmed enhanced biosynthesis of asparagine in M cells in comparison to C1, from both glucose and glutamine (Figure 4N and O). This is consistent with the observed significant increase in asparagine synthetase (ASNS) abundance in M cells in comparison to C1 in our proteomics analysis (Figure 4E). Together, these data demonstrate aberrant amino acid metabolism across colorectal tumour development and in particular highlight a potential role for the asparagine biosynthesis pathway in promoting CRC progression.

### *ASNS* expression is increased in colorectal cancer and supports tumour cell phenotype

As ASN biosynthesis (summarised in Figure 5A) was increased across CRC progression, and ASNS abundance was elevated in M cells in comparison to C1, we decided to assess ASNS abundance across our cell line series. We confirmed increased expression of ASNS in late-stage M cells and revealed higher abundance in the SB and 10C cells in comparison to C1 cells by immunoblot (Figure 5B). Consistent with this, using The Cancer Genome Atlas (TCGA) data we found that *ASNS* expression was significantly elevated in human colorectal tumour (2.64-fold) and metastatic (2.22-fold) tissue in comparison to normal (Figure 5C). High levels of *ASNS* expression were also associated with significantly poorer overall survival in CRC patients using two separate publicly available CRC datasets (GSE17536; Figure 5D; n=174; HR 1.63; p=0.013 and GSE29621; Supplementary Figure S5A; n=65; HR 2.64; p=0.004).

**Figure 5.**
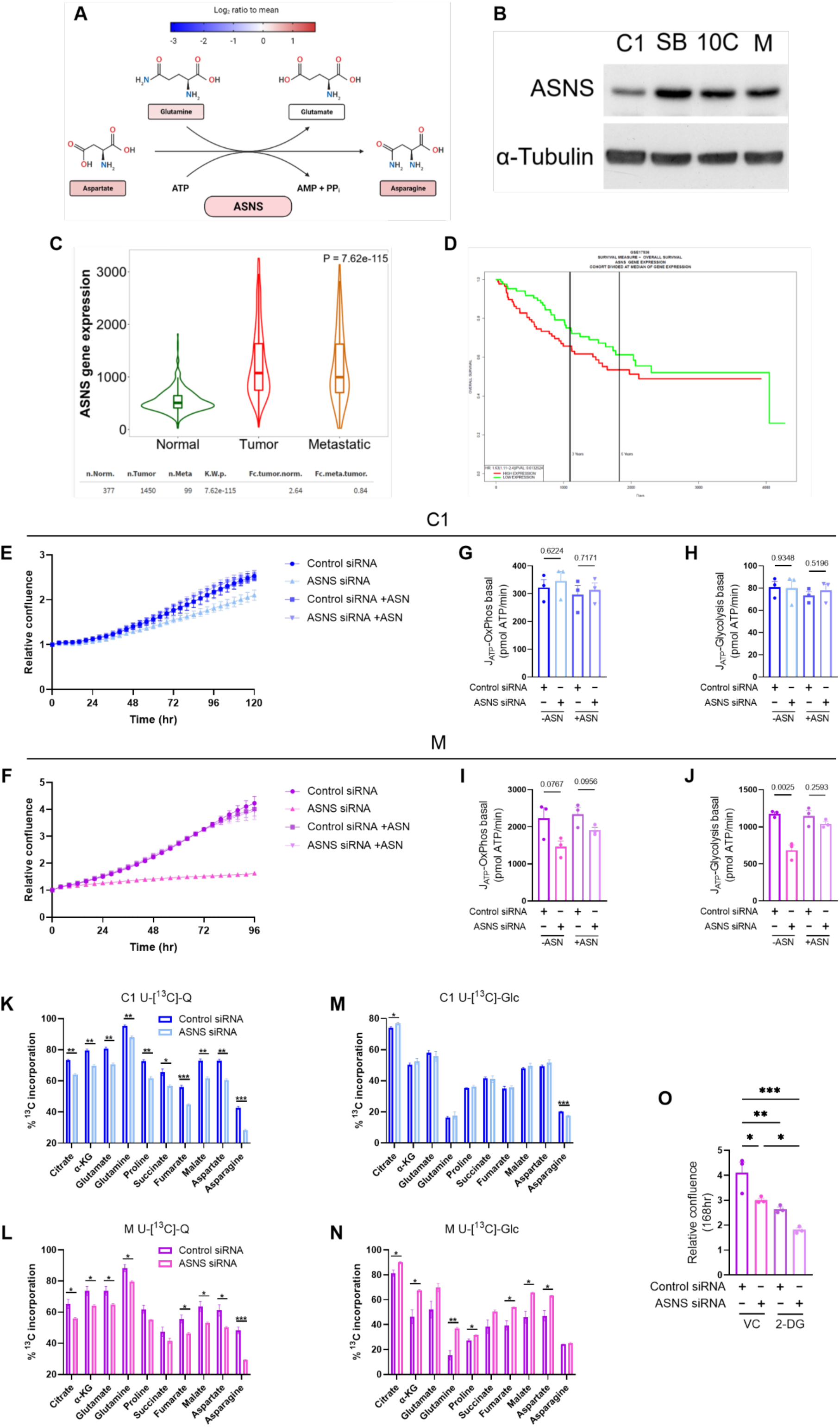
***ASNS* expression is increased in colorectal cancer and supports tumour cell proliferation through ASN synthesis** (A) Schematic of the asparagine biosynthesis pathway. Colour of amino acids (rectangles) represents their relative abundance in M cells (log_2_ ratio to mean for each amino acid) using data from heatmap in Figure 4D. Colour of asparagine synthetase (ASNS) represents significant upregulation of ASNS in M/C1 using data from Figure 4E. Figure created using BioRender.com. (B) Immunoblot of ASNS abundance in cell lines indicated. α-Tubulin serves as loading control. Representative of three independent experiments. (C) Violin plot of *ASNS* expression in normal (n=377), tumour (n=1450) and metastatic (n=99) human colorectal tissue using TNMplot (22) and publicly available gene chip data from The Cancer Genome Atlas (TCGA). *ASNS* expression was significantly increased in colorectal tumours (2.64-fold; Dunn’s p=1.84×10^-116^) and metastases (2.22-fold; Dunn’s p=3.38×10^-2^) compared to normal colonic tissue. (D) Overall survival analysis in relation to *ASNS* expression from GSE17536 (23). Cohort divided at median of *ASNS* expression. n=174; HR 1.63; p=0.013. (E & F) Proliferation of C1 (E) and M (F) cells following transfection with non-targeting control (Control) or *ASNS*-targeting siRNA, with or without 0.1 mM asparagine (ASN) supplementation. Basal ATP generation via OxPhos in C1 (G) and M (I) cells and basal ATP generation via glycolysis in C1 (H) and M (J) cells following transfection with non-targeting control (Control) or *ASNS*-targeting siRNA, with or without 0.1 mM ASN supplementation. (K-N) Stable isotope labelling (SIL) using uniformly labelled U-[^13^C]-Q (K & L) or U-[^13^C]-Glc (M & N). Data expressed as percentage ^13^C incorporation into TCA cycle intermediates and associated amino acids in C1 (K & M) and M (L & N) cells. (O) Proliferation of M cells following transfection with non-targeting control (Control) or *ASNS*-targeting siRNA, with vehicle control (VC) or 10 mM 2-Deoxy-D-Glucose (2-DG). Assay performed as in (E & F). (E-O) Data are represented as mean ± SEM of three independent cultures. (G-N) Student’s t-test. (O) Tukey’s multiple comparisons test *p<0.05; **p<0.01; ***p<0.001.

Given these findings, we next set out to investigate the role of ASNS in colorectal tumour cell phenotype. Proliferation was compared using an Incucyte Zoom live cell imaging system in early C1 cells and late M cells following *ASNS* knockdown using RNAi. Parallel samples from these experiments were used to confirm *ASNS* knockdown (see Figure 6A-D). We found that proliferation of C1 cells was only moderately impaired by *ASNS* knockdown (17% reduction in confluence versus control siRNA at 120 hours; Figure 5E; Supplementary Figure S5B), whereas M cell proliferation was almost entirely blunted following suppression of *ASNS* expression (61% decrease in confluence versus control siRNA at 96 hours; Figure 5F; Supplementary Figure S5C). The phenotype was rescued by exogenous ASN, demonstrating the growth promoting effects of ASNS are attributable to its role in ASN synthesis (Figure 5E and F; Supplementary Figure S5B and C). Inhibition of proliferation via *ASNS* knockdown appears to be cytostatic rather than cytotoxic, as there were no differences in apoptosis detected by caspase 3/7 staining (Supplementary Figure S5D and E).

**Figure 6.**
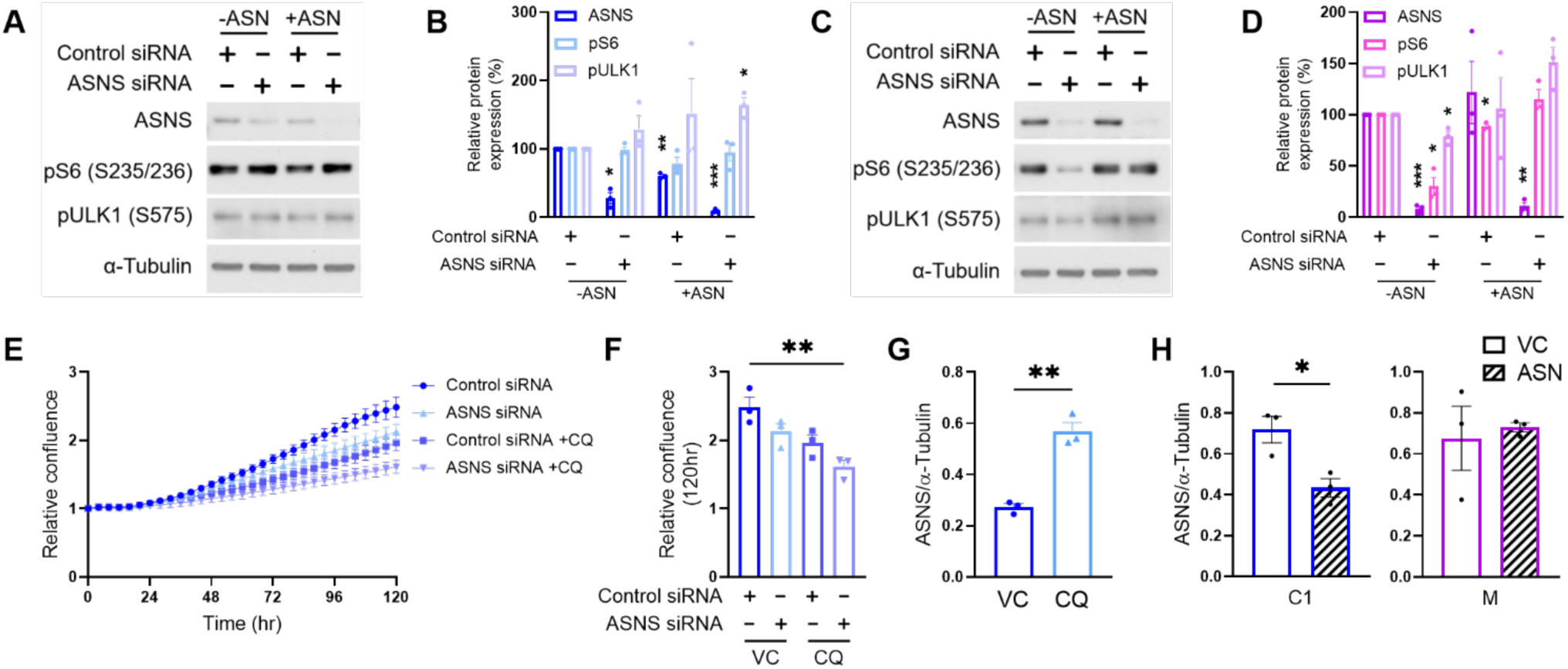
**ASNS activity is required for maintaining mTORC1 signalling in adenocarcinoma cells but is dispensable in early adenomas** (A-D) Analysis of mTORC1 signalling following siRNA-mediated ASNS suppression. Immunoblots of ASNS, phosphorylated ribosomal S6 protein (S235/236) and phosphorylated ULK1 (S575) expression following transfection with non-targeting control or *ASNS*-targeting siRNA, with or without 0.1 mM ASN supplementation in C1 (A) and M (C) cells. α-Tubulin serves as loading control. Quantification of three independent cultures by densitometry using Image J in C1 (B) and M (D) cells. (E & F) Proliferation of C1 cells following transfection with non-targeting control or *ASNS*- targeting siRNA, with or without 10 µM chloroquine (CQ). Water used as vehicle control (VC). (G) Relative abundance of ASNS protein (ASNS/α-Tubulin) in C1 cells following 10 µM CQ addition. (H) Relative abundance of ASNS protein (ASNS/α-Tubulin) following 0.1 mM ASN supplementation in cell lines indicated. (G & H) Quantification of three independent cultures by densitometry using Image J. (A-H) Data are represented as mean ± SEM of three independent cultures. (B & D) One sample t-test. (F) Tukey’s multiple comparisons test. (G & H) Student’s t-test. *p<0.05; **p<0.01; ***p<0.001.

To explore the metabolic implications of *ASNS* knockdown in the relatively insensitive C1 adenoma cells in comparison to the sensitive M adenocarcinoma cells, we carried out extracellular flux analysis. Consistent with the minimal effect of *ASNS* knockdown on proliferation, we observed no measurable differences in OCR or ECAR following *ASNS* knockdown in the C1 adenoma cells (Figure 5G and H; Supplementary Figure S5F- H). However, *ASNS* suppression in the M adenocarcinoma cells showed reduced basal OCR and basal and max ECAR, which was rescued by ASN addition (Figure 5I and J; Supplementary Figure S5I-K). These data indicate ASNS activity supports OxPhos and glycolysis in late-stage colorectal carcinoma cells, but this association is absent in early adenoma cells.

To explore this further, we carried out SIL using U-[^13^C]-Glc and U-[^13^C]-Q in C1 and M cells transfected with control or *ASNS* targeting siRNA. Incorporation of glutamine is reduced in TCA cycle intermediates in both the C1 (Figure 5K) and M cells (Figure 5L) following *ASNS* suppression. However, the cell lines diverge regarding glucose incorporation into the TCA cycle. There was largely no major difference in glucose labelling in TCA cycle intermediates upon *ASNS* knockdown in C1 adenoma cells (Figure 5M); however, glucose incorporation was increased in almost all TCA cycle intermediates in the M adenocarcinoma cells upon *ASNS* suppression (Figure 5N). These data suggest that M cells compensate for reduced glutamine anaplerosis by diverting glucose into the TCA cycle away from lactate production (hence reduced ECAR with *ASNS* knockdown – Figure 5J), in an insufficient effort to maintain OxPhos. To target this metabolic fragility unique to the late-stage adenocarcinoma cells, we used 2-deoxy-D-glucose (2-DG) – a glucose analogue and glycolysis inhibitor (24-26) – as a means of preventing compensatory glucose shuttling and further inhibiting tumour cell proliferation mediated by *ASNS* knockdown (Figure S5L). We found 2-DG in combination with *ASNS* suppression led to a further decrease in M adenocarcinoma cell proliferation (Figure 5O; Supplementary Figure S5M and N). These data indicate that using 2-DG to target compensatory glucose shuttling enhances the anti-proliferative impact of *ASNS* knockdown in later stage adenocarcinoma cells.

### Regulation of asparagine availability is a node of metabolic vulnerability in later stage colorectal adenocarcinoma cells

The role of ASN in mTORC1 activation has recently been established (27, 28), and mTORC1 signalling is well known to promote cellular growth and proliferation (29). We therefore analysed phosphorylation of mTORC1 targets following siRNA-mediated suppression of *ASNS* in the early C1 cells vs the later stage M cells. In the C1 cells, immunoblot analyses revealed that *ASNS* knockdown did not impact phosphorylation of mTORC1 signalling targets; S6 ribosomal protein and ULK1 (Figure 6A and B). By contrast, in M cells, mTORC1 activity appeared tightly coupled to *ASNS* expression, indicated by significantly reduced S6 and ULK1 phosphorylation following *ASNS* suppression, which is reversed following the addition of asparagine (Figure 6C and D).

An explanation for this could be that C1 cells remain able to source ASN in the absence of ASNS activity, which may also explain their insensitivity to *ASNS* knockdown in proliferation assays (Figure 5E). We hypothesised that adenoma cells may be resistant to *ASNS* knockdown and able to maintain mTORC1 activity as they are able to scavenge ASN through autophagy (30, 31). In an attempt to sensitise C1 cells to *ASNS* knockdown, we treated them with the autophagy inhibitor chloroquine (CQ) and analysed cell proliferation over 120 hours. Critically, CQ-mediated inhibition of autophagy sensitised the adenoma cells to *ASNS* knockdown, significantly reducing proliferation in comparison to control cells (Figure 6E and F; Supplementary Figure S6A).

Interestingly, we observed that CQ upregulated abundance of ASNS in adenoma cells (Figure 6G; Supplementary Figure S6A), presumably a mechanism for attempting to restore intracellular ASN levels, suggesting ASN biosynthesis and autophagy are intrinsically linked in adenoma cells. Moreover, further analysis of immunoblot data (Figure 6A-D) revealed significantly reduced ASNS abundance in the C1 adenoma cells, but not the M adenocarcinoma cells, following ASN supplementation (Figure 6H). These observations suggest a feedback loop present in early adenoma cells (to sense and regulate ASN levels) to maintain a check on proliferation, that is lost in the late-stage cells. M adenocarcinoma cells require ASN yet have lost the ability to sense and regulate its levels using autophagy. Together, our findings suggest ASNS is a node of metabolic vulnerability in later stage colorectal adenocarcinoma cells.

## Discussion

The aim of this study was to understand how metabolism changes across CRC development and to identify tractable nodes of metabolic vulnerability with potential for CRC treatment. To do this we used a unique *in vitro* model of CRC progression allowing us to study metabolic changes at the very earliest, right through to the later, more aggressive stages of human colorectal tumour progression. Using this model allowed us to characterise several distinct metabolic changes that occur across progression leading to the identification of the dependency of late-stage cells on ASNS (Figure 7A).

**Figure 7.**
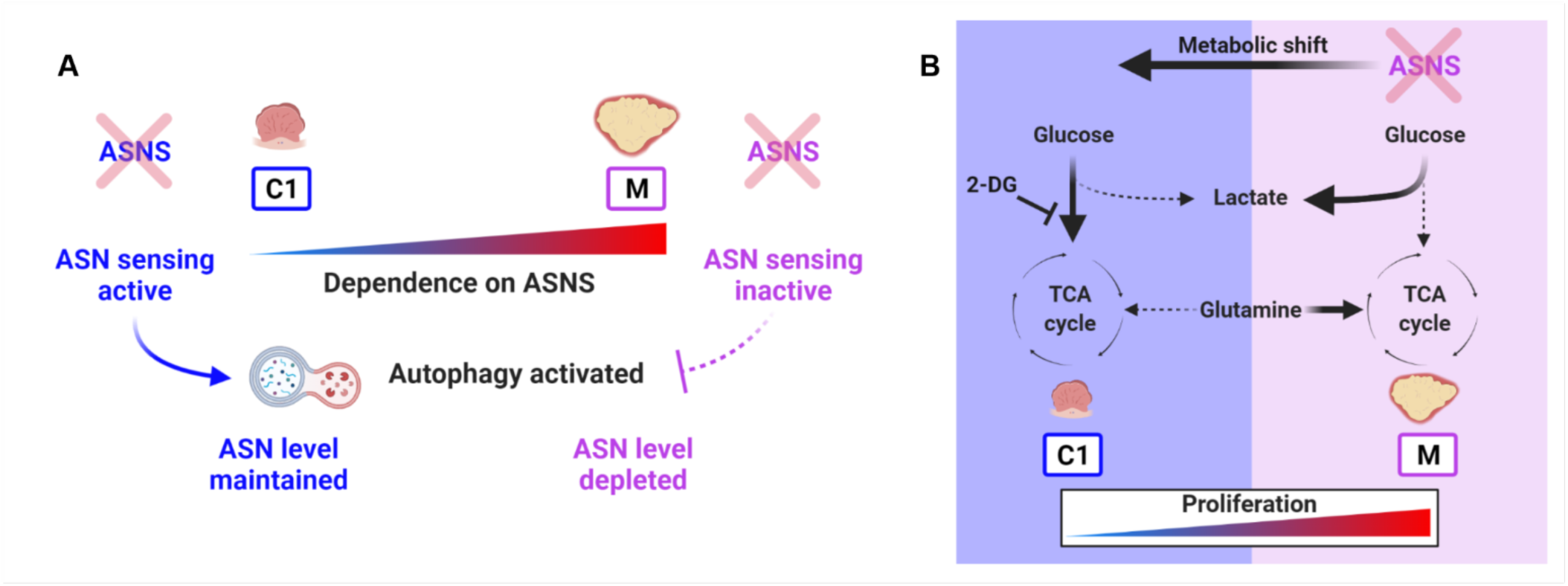
**ASNS is a node of metabolic vulnerability in later stage colorectal adenocarcinoma cells** (A) Late-stage adenocarcinoma cells (M) have become addicted to ASN but have an inactive ASN-sensing mechanism. This renders them unable to sense and react to levels of ASN via autophagy activation, leaving them with insufficient ASN to maintain their high proliferative rate. Thus, M cells are vulnerable to ASNS depletion in the absence of extracellular ASN. Early adenoma cells (C1) can sense high or low ASN levels and regulate expression of *ASNS* accordingly to keep a check on cellular proliferation, or in the event of ASNS loss can activate autophagy to maintain intracellular ASN level (Figure 6). (B) Targeting ASNS in late adenocarcinoma (M) cells using siRNA reduces glutamine-derived carbon flux into the TCA cycle. This causes compensatory influx of glucose-derived carbon into the TCA cycle, diverting glucose carbon away from lactate production (Figure 5). This shifts the metabolic and proliferative phenotype of M cells back towards that of the early adenoma (C1) cells. Targeting this compensatory glucose shuttling into the TCA cycle with 2-DG further sensitises M cells to ASNS suppression (Figure 5O). Figure created using BioRender.com.

Importantly, our model has allowed us to explore the dynamic metabolic changes that characterise the very earliest stages of CRC development. We show that colorectal tumour cells switch early to a glycolytic phenotype (from C1 to SB; Figure 2F), elevating lactate production but maintaining oxidative metabolism. This transition is maintained throughout the series (SB to 10C to M). Our data are consistent with studies of colorectal adenomas *in vivo*, where increased consumption of glucose (13, 32, 33) and inositol (13) were observed, supporting the idea that a glycolytic switch occurs early in tumorigenesis and validating our progression model used in this study. Our SIL data reveal that the early glycolytic switch is accompanied by enhanced glutaminolysis, which is also sustained through progression (Figure 2F). Thus, at this early transition (C1 to SB), increased glutamine-anaplerosis maintains the TCA cycle, compensating for the deficit in glucose carbon which is diverted instead towards lactate production. Increased glutaminolysis is an established feature of colorectal carcinoma cells (34-38), however, our data suggest this is an early reprogramming event, initiated in the pre-malignant stages of CRC. These findings could have important implications for prevention strategies at an early, pre-cancerous stage. Our data suggest treatment with glutaminase inhibitors such as CB-839 (39, 40) could be efficacious at this pre-malignant stage with potential to provide protection against recurrence, generation of new polyps or progression to malignancy, although further studies will be required.

Mutations that activate the WNT pathway are extremely common driver events in CRC, with mutation of the *APC* tumour suppressor gene the most frequent initiating event (41). In our model, the C1 cells were derived from a polyp from a familial adenomatous polyposis (FAP) patient. These patients are born with a germline mutation affecting one *APC* allele, spontaneous loss of the second wild-type allele leads to enhanced WNT signalling and adenoma formation at an early age (42). It is worth noting that aberrant WNT pathway activation has been associated with some of the metabolic phenotypes we observe at the early stages of tumour progression in our model. For example, enhanced glycolysis has been reported through WNT-mediated regulation of PDK1 (43) and pyruvate carboxylase (44) expression, and WNT target c-MYC has been shown to promote glutaminolysis (45, 46). It may be that genetic events that deregulate WNT signalling could underlie some of the metabolic phenotypes we observe. However, as our series of cell lines were initially derived from a polyp already harbouring aberrant WNT activation due to loss of both wild-type *APC* alleles, it is more likely that alternative underlying driver mutations account for changes in metabolism we see in our model.

We found that oxidative metabolism was maintained throughout CRC progression and was enhanced in the most progressed adenocarcinoma cells (M), supporting its consideration for therapeutic intervention (39, 47, 48). It has been shown that CRC cells consume more oxygen and contain more mitochondria than surrounding normal colonic tissue *in vivo* (49). Transmission electron microscopy data did not indicate any significant differences in mitochondrial number throughout progression (Supplementary Figure S1), although our analyses cannot rule out differences in cristae morphology.

Despite demonstrating metabolic plasticity with respect to nutrient source (supporting their ability to proliferate in nutrient-poor environments); importantly, we found that adenocarcinoma cells were highly dependent on *de novo* ASN biosynthesis. This is supported by clinical data showing *ASNS* expression is elevated in colorectal tumour and metastatic tissue in comparison to normal and is associated with poorer overall survival. Consistent with this, high levels of ASN have previously been found to promote breast cancer metastases, with enhanced *ASNS* expression in primary tumours strongly correlating with metastatic potential (50). In our model, enhanced ASNS abundance in the more progressed cells may be associated with activating mutations in the KRAS oncogene (14), which has previously been linked to increased *ASNS* expression in CRC cells (51).

Loss of *ASNS* expression in late-stage cells blocked proliferation, while only minimally disadvantaging the early-stage cells. The phenotype was rescued with exogenous ASN demonstrating that it is reduced supply of the amino acid that impacts the cells rather than the physical presence of the enzyme or another by-product of its activity. Previous studies in CRC cells have shown that manipulation of ASN/ASNS only results in growth arrest in the context of glutamine depletion (30, 51). However, we observe a strong phenotype in glutamine replete conditions suggesting our cells are dependent on ASN regardless of glutamine availability.

Another important discovery from our study is the ASN-mediated potentiation of glycolytic metabolism in human CRC. At the time of writing, we found one other study - by Xu *et al.* - reporting that ASNS/ASN manipulation affects glycolysis in the context of thermogenesis in brown adipose tissue in mice (52). We also show that *ASNS* suppression reduces glutamine-derived carbon flux into the TCA cycle. Interestingly, these metabolic changes demonstrate a shift back towards the metabolic phenotype characterising the early-stage cells (Figure 7B). This shift to glucose utilisation sensitised cells to 2-DG, targeting the compensatory glucose carbon shuttling into the TCA cycle. These data support the growing body of evidence highlighting the importance of combination therapies when targeting metabolic vulnerabilities in cancer (7, 39, 53).

In agreement with the Xu study, and others by Krall *et al.* (27, 28), our data show that ASN promotes mTORC1 signalling. However, we show that dependence on *ASNS* expression for maintenance of mTORC1 activity in colorectal tumour cells is very much dependent on stage of progression. We found that mTORC1 signalling was inhibited in M cells upon ASNS loss (this was reversed by ASN supplementation), but not in C1 cells. This led us to hypothesise that C1 cells were able to source ASN through an ASNS-independent mechanism, such as autophagy, which has been previously observed in CRC cells (30, 31). We found that the autophagy inhibitor CQ sensitised early adenoma cells to *ASNS* suppression, further inhibiting proliferation of the early-stage cells. We therefore identified a feedback mechanism for sensing and regulating ASN levels that is present in early adenoma cells, allowing them to maintain a check on proliferation - that is lost in late adenocarcinoma cells - leaving these cells vulnerable to manipulation of the ASN/ASNS axis.

ASNS therefore represents an exciting potential target for CRC therapy. But it is important to remember that targeting ASNS will only be effective in the context of asparaginase treatment or upon blockade of ASN uptake. Asparaginase has been used to treat acute lymphoblastic leukaemia in the clinic for decades by reducing levels of circulating ASN (54). However, these cancers eventually develop resistance by increasing *de novo* ASN synthesis through upregulation of *ASNS* expression (55, 56). Interestingly, asparaginase has been shown to be effective in treating CRC with mutations that inhibit GSK3α – such as those caused by R-spondin fusions – but was ineffective in APC and β-catenin mutant CRCs, which represent the vast majority of cases. This same study demonstrated that pharmacological inhibition of GSK3α could sensitise APC and β-catenin mutant CRCs to asparaginase by preventing the release of ASN through GSK3α-mediated protein degradation (31). Our results agree that preventing access to ASN in APC mutant cells – both at the adenoma and carcinoma stage – may be critical when targeting ASNS for CRC therapy. Specific ASNS inhibitors are currently in development (57) and may offer effective future treatment strategies for CRC, and indeed other cancers, in combination with asparaginase and/or autophagy inhibition. We have therefore demonstrated the strength of our model for the identification of metabolic dependencies and potential therapeutic strategies, exemplified by the identification of ASNS as a node of metabolic vulnerability in CRC.

## Acknowledgements

We thank D. Avizonis and L. Choinière from McGill University Metabolomics Core Facility, Kate Heesom and Phil Lewis from University of Bristol Proteomics Facility and the Wolfson Bioimaging Facility at the University of Bristol. EEV, DNL and CJB are supported by Diabetes UK (17/0005587) and the World Cancer Research Fund (WCRF UK), as part of the World Cancer Research Fund International grant program (IIG_2019_2009). SG is funded by Above and Beyond Charity and BA is supported by MRC grant MR/R02149X/1. EEV and ACW are supported by the CRUK Integrative Cancer Epidemiology Programme (C18281/A29019). EEV and CJB work in a unit funded by the UK Medical Research Council (MC_UU_00011/1 & MC_UU_00011/4) and the University of Bristol.

## Materials and Methods

### Cell lines and culture

The colorectal adenoma-derived cell line PC/AA/C1 (C1), and the transformed adenoma-derived cell lines PC/AA/C1/SB (SB), PC/AA/C1/SB10 (10C) and PC/AA/C1/SB10/M (M) were generated in the Paraskeva lab, their derivation and characterisation have been previously described (14, 58). Growth medium was Dulbecco’s modified Eagle’s medium (DMEM) (Gibco; Thermo Fisher Scientific, MA, USA) supplemented with 20% foetal bovine serum (FBS), 1 µg/mL hydrocortisone sodium succinate (Sigma), 0.2 U/mL insulin (Sigma), 4mM glutamine (Gibco), 100 U/mL penicillin and 100 μg/mL streptomycin (Gibco). All cell lines were routinely assessed for microbial contamination (including mycoplasma). Stocks were securely catalogued and stored, and passage numbers strictly adhered to prevent phenotypic drift.

### Immunoblotting

Whole-cell lysates were prepared in situ and analysed by western blotting as previously described (14) using antibodies to the following: ASNS (20843, Cell Signaling Technology; 1:1000), phospho-S6 ribosomal protein S235/236 (4858; Cell Signaling Technology; 1:2000), phospho-ULK1 S757 (6888; Cell Signaling Technology; 1:1000), Total OXPHOS Human Antibody Cocktail (Ab110411; Abcam, Cambridge, UK; 1:200), α-tubulin (T9026; Sigma; 1:10000). Where indicated, three independent blots were quantified by densitometry analysis using ImageJ software version 1.51r.

### RNA interference (RNAi)

Cells were reverse transfected in Opti-MEM (Gibco) with small interfering RNA (siRNA) using Lipofectamine RNAiMAX (Invitrogen, Thermo Fisher Scientific, MA, USA) according to manufacturer’s instructions. Smartpool (pool of four distinct siRNA sequences) negative control siRNA, and Smartpool *ASNS*-targeting siRNA (Dharmacon, Horizon Discovery, Cambridge, UK; 50 nM) were used in this study. Cells were incubated overnight at 37 °C before medium changing and supplementation with asparagine (ASN; A4159; Sigma; 0.1 mM) chloroquine (CQ; C6628; Sigma; 10 µM), or 2-Deoxy-D-Glucose (2-DG; D8375; Sigma; 10 mM), as stated.

### Extracellular flux analyses

Mitochondrial stress test assays were carried out using the Seahorse XFe96 Analyzer (Agilent Technologies) and data were acquired using Seahorse Wave software v2.6 (Agilent Technologies). Injections of drugs during the assay were at timepoints stated in figures. Injections were of oligomycin (75351; Sigma; 2 µM), FCCP (C2920; Sigma; 1 µM), antimycin A (A8674; Sigma; 1 µM), rotenone (R8875; Sigma; 1 µM) and monensin (M5273; Sigma; 20 µM). Raw ECAR and OCR measurements taken by the instrument from individual wells were normalised using crystal violet staining of plates immediately upon completion of the assay as follows: cells were fixed in 4% PFA and stained using 0.5% crystal violet (Sigma) before solubilisation in 2% SDS (Severn Biotech, Worcs, UK) and subsequent OD^595^ measurements were obtained using an iMark microplate reader (Bio-Rad). Data were normalised by dividing raw OCR/ECAR measurement by OD^595^ for each well of the culture plate. Bioenergetics analyses were carried out by taking normalised OCR/ECAR measurements and applying them to a bioenergetics spreadsheet developed by Ma *et al.* based on ATP calculations from Mookerjee *et al.* (16, 17) to generate J-ATP production data.

#### Analysis of cell line progression model

Cells were seeded in regular growth medium (20% DMEM) into XF96 V3 PS Cell Culture Microplates (Agilent Technologies) then medium changed the following day. Following 72 hours of further culture, cells were washed once then fed Seahorse XF DMEM assay media (103575; Agilent Technologies) supplemented with 10 mM glucose (103577; Agilent Technologies), 2 mM glutamine (103579; Agilent Technologies) and 1 mM sodium pyruvate (103578; Agilent Technologies) for 1 hour prior to assay.

#### Nutrient restriction analyses

Cells were seeded in regular 20% DMEM growth media into XF96 V3 PS Cell Culture Microplates then medium changed the following day. Following 72 hours of further culture, cells were washed once then fed Seahorse XF assay media supplemented with either 10 mM glucose (for glutamine restriction) or 2mM glutamine (for glucose restriction) for 2 hours prior to assay.

#### ASNS knockdown analyses

*ASNS* expression was suppressed using RNAi (as stated previously) and cells were seeded into XF96 V3 PS Cell Culture Microplates then medium changed the following day with 20% DMEM or 20% DMEM supplemented with 0.1 mM ASN. Following 72 hours of further culture, cells were washed once then fed Seahorse XF assay media supplemented with 10 mM glucose, 2 mM glutamine and 1 mM sodium pyruvate for 1 hour prior to assay.

### Stable Isotope Labelling (SIL)

U-[^13^C]-Glc labelling media consisted of DMEM supplemented with 10 mM D-glucose (U-^13^C6; CLM-1396; Cambridge Isotope Laboratories), 2 mM glutamine, 100 U/mL penicillin and 100 μg/mL streptomycin, 0.4 mM phenylalanine (78019; Sigma) and 10% dialysed FBS (Gibco). U-[^13^C]-Q labelling media consisted of DMEM supplemented with 2 mM L-glutamine (U-^13^C5; CLM-1822; Cambridge Isotope Laboratories), 10 mM glucose, 100 U/mL penicillin and 100 μg/mL streptomycin and 10% dialysed FBS. Following labelling, cells were washed twice with saline and lysed with 0.8 mL methanol, sonicated for 3×30 second pulses, and dried using a speed vacuum. Gas chromatography coupled to mass spectrometry (GC/MS) was performed at McGill University using previously detailed methods (59) and MIDs were derived using an algorithm developed at McGill University (60). Relative metabolite levels were normalised to cell number using counts from duplicate plates with Cytosmart cell counter (Corning).

#### SIL of cell line progression model

Cells were seeded in regular 20% DMEM into 6cm plates for 72 hours. Cells were washed twice with PBS prior to pulse with U-^13^C-glucose or U-^13^C-glutamine labelling media for 24 hours. Cells were then lysed and intracellular metabolites analysed by GC/MS as described above.

#### SIL following ASNS knockdown

*ASNS* expression was suppressed using RNAi (as stated previously) and cells were seeded into 6cm plates (Corning). Plates were medium changed with 20% DMEM the following day and cultured for a further 48 hours. Cells were washed twice with PBS prior to pulse with U-^13^C-glucose or U-^13^C-glutamine labelling media for 24 hours. Cells were then lysed and intracellular metabolites analysed by GC/MS as described above.

### Transmission electron microscopy (TEM)

Cells were grown in T25 flasks (Corning, NY, USA) for 48 hours, washed twice with PBS and fixed with glutaraldehyde. Samples were prepared as detailed previously (61) by the Wolfson Bioimaging Facility, University of Bristol. Images of cell lines in the progression model obtained via TEM were imported into Adobe Photoshop for analyses of mitochondrial number and morphology. Each mitochondrion within a cell was analysed for area, length and roundness. A minimum of 10 cells per cell line were analysed.

### Nutrient restriction proliferation assays

Cells were seeded into 96 well plates (Corning) in full 20% DMEM for 72 hours. Cells were then washed twice with PBS and media containing reduced glucose or glutamine was added. Reduced glucose media consisted of DMEM supplemented with 4mM glucose (Agilent Technologies), 2 mM glutamine, 100 U/mL penicillin and 100 μg/mL streptomycinand 10% dialysed FBS (Gibco). Reduced glutamine media consisted of DMEM supplemented with 0.5 mM glutamine (Agilent Technologies), 10 mM glucose (Agilent Technologies), 100 U/mL penicillin and 100 μg/mL streptomycin and 10% dialysed FBS (Gibco). Plates of cells were subsequently fixed at day 0 (day of treatment), day 3, day 5 and day 7. Cells were fixed in 4% PFA and stained using 0.5% crystal violet (Sigma) before solubilisation in 2% SDS (Severn Biotech, Worcs, UK) and subsequent OD^595^ measurements were obtained using an iMark microplate reader (Bio-Rad). Data are expressed as OD^595^ of selected timepoint relative to OD^595^ at day 0.

### *ASNS* knockdown proliferation assays

*ASNS* expression was suppressed using RNAi (as stated previously) and cells were seeded into 96 well plates. Cells were medium changed the following day with 20% DMEM or 20% DMEM supplemented with 0.1 mM ASN, 10 mM 2-DG or 10 µM CQ, as indicated in text and figures. Plates were immediately incubated in IncuCyte ZOOM live cell imaging system. Phase confluence (%) was measured every four hours for times indicated in graphs. IncuCyte system scanned and took images at four different fields in each well. For each individual well, phase confluence measurements taken at each timepoint were measured relative to the initial measurement (0 hours) and data were expressed as relative confluence measurements. For ASNS knockdown 2-DG experiments, CellEvent caspase-3/7 green detection reagent (C10423; Invitrogen; 1:1000) was used to measure apoptosis. For each individual well, green object count (1/mm^2^) was normalised to relative confluence at each timepoint, and results expressed as relative apoptosis.

### Tandem Mass Tag (TMT) proteomic analysis

Cells were seeded in 20% DMEM and cultured in T25 flasks for 48 hours, then medium changed and cultured for a further 24 hours. Whole cell lysates were prepared from cells as previously described (14). Lysates were submitted to the proteomics facility at the University of Bristol where TMT proteomic analysis was carried out, as previously described (62). Briefly, samples were labelled with TMT nine plex reagents according to the manufacturer’s protocol (Thermo Fisher Scientific, Loughborough, LE11 5RG, UK), and the labelled samples were pooled prior to fractionation by high pH reversed-phase chromatography using an Ultimate 3000 liquid chromatography system (Thermo Fisher Scientific). Fractions were evaporated to dryness and resuspended in 1% formic acid prior to analysis by nano-LC MSMS using an Orbitrap Fusion Tribrid mass spectrometer (Thermo Scientific). All spectra were acquired using an Orbitrap Fusion Tribrid mass spectrometer controlled by Xcalibur software (Thermo Scientific). The raw data files were processed and quantified using Proteome Discoverer software (Thermo Scientific) and searched against the UniProt Human database. All peptide data were filtered to satisfy false discovery rate (FDR) of 5%. Significantly regulated proteins (FDR<5%, -log10 p-value>1.3 and log2 fold change+/-0.48) were selected for further pathway enrichment analyses using Gene Ontology (GO) (63, 64) and Kyoto Encyclopedia of Genes and Genomes (KEGG) (65).

### *ASNS* expression in tumour, normal and metastatic colon tissue

Analysis of *ASNS* expression was performed in normal (n=377), tumour (n=1450) and metastatic (n=99) human colorectal tissue using TNMplot (22) with publicly available gene chip data from Gene Expression Omnibus (GEO) (66) and The Cancer Genome Atlas (TCGA; https://www.cancer.gov/tcga).

### Survival analysis

Survival analyses in relation to *ASNS* expression were generated using PROGgeneV2 (23) and publicly available CRC datasets (GSE17536; GSE29621) from GEO (66).

### Statistical analysis

Statistical analyses were performed in GraphPad Prism using One-way ANOVA with post-hoc Tukey’s multiple comparison’s test, Student’s t-test or one sample t-test, as stated. Significance was expressed as \**p*<0.05; \*\**p*<0.01; \*\*\**p*<0.001; \*\*\*\**p*<0.0001. Results are expressed as mean ± SEM of three independent cultures, unless otherwise stated.

