## Supplementary Figures for "Identifying targetable metabolic dependencies across colorectal cancer progression"

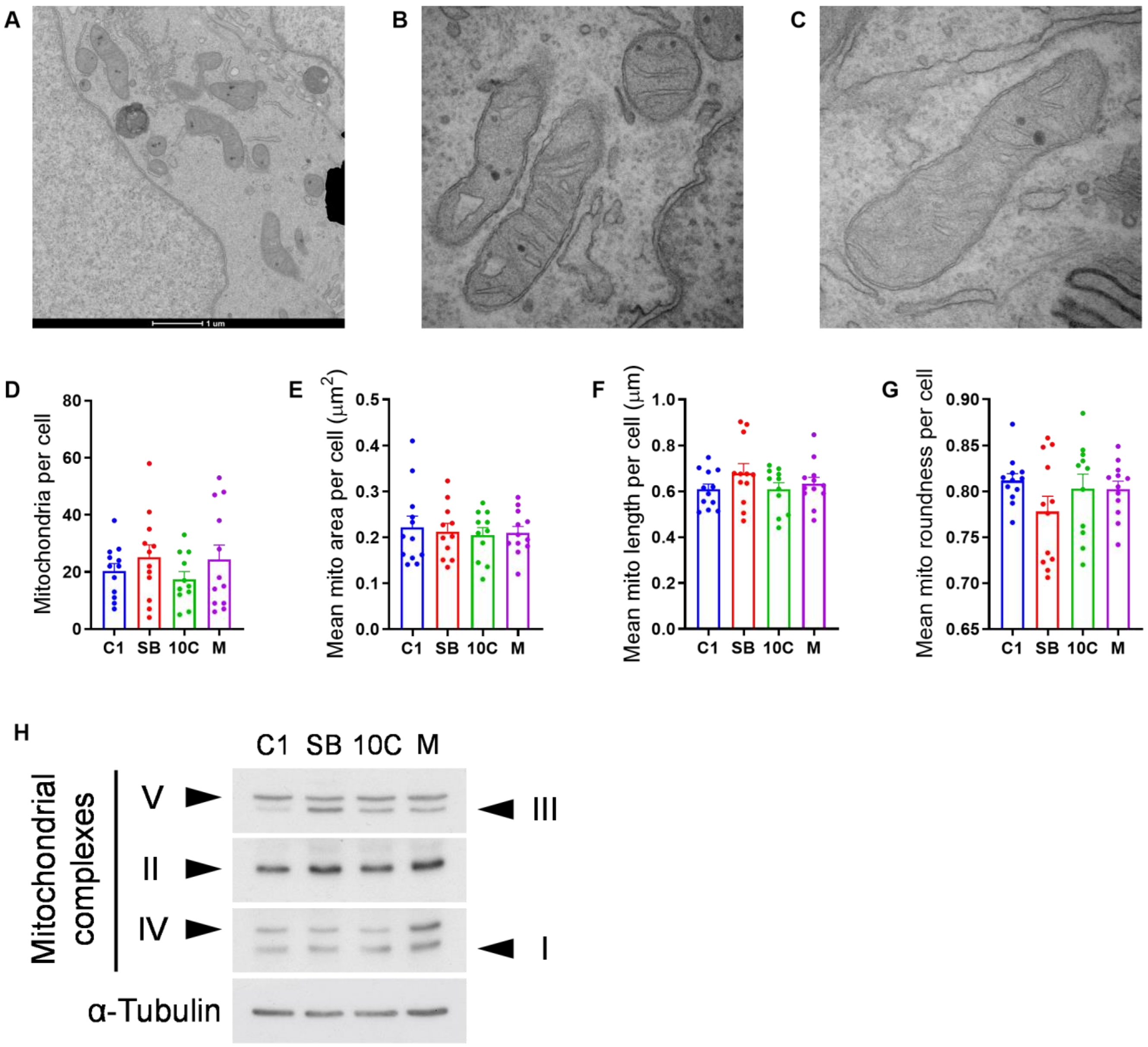


**Supplementary Figure S1. Mitochondrial morphology analyses**

(A-C) Example transmission electron microscopy (TEM) images of mitochondria. (A) 6800x (B & C) 30000x magnification. (D-G) Measurements of mean mitochondrial number (D), area (E), length (F) and roundness (G) per cell using Adobe Photoshop. Data are represented as mean ± SEM of a minimum of 10 cells per cell line. (H) Immunoblot of mitochondrial complex I – V expression in cell lines indicated. α-Tubulin serves as loading control. Representative of three independent experiments.

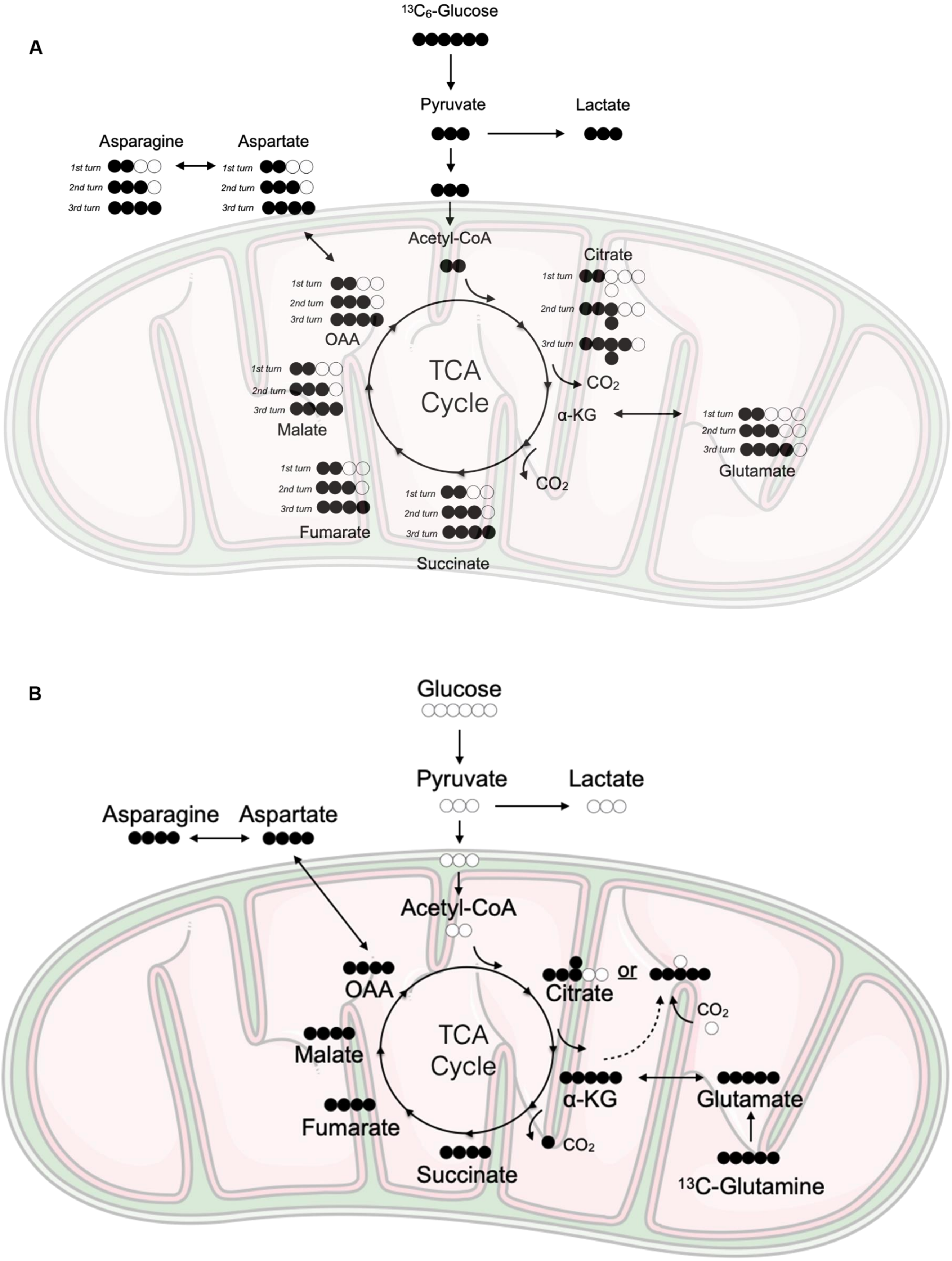


**Supplementary Figure S2. Stable isotope labelling schematics**

(A & B) Stable isotope labelling schematics depicting the generation of different isotopologues of indicated metabolites when using uniformly labelled $^{13}$C-glucose (U-[$^{13}$C]-Glc) (A) or 13C-glutamine (U-[$^{13}$C]-Q) (B). Black circles represent $^{13}$C atoms. White circles represent $^{12}$C atoms.

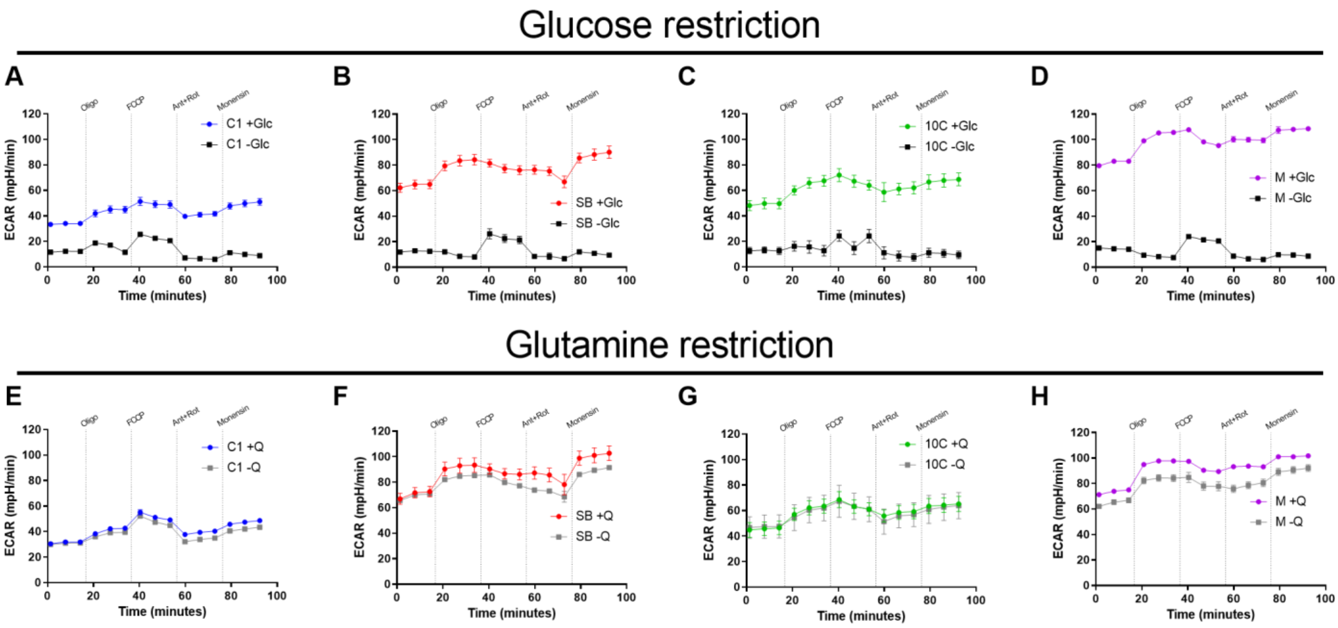


**Supplementary Figure S3. Extracellular acidification rate during nutrient restriction**

(A-D) Extracellular acidification rate (ECAR) traces in C1 (A), SB (B), 10C (C) and M (D) cells in presence (+Glc; 10 mM) or absence (-Glc; 0 mM) of glucose. (E-H) ECAR traces in C1 (E), SB (F), 10C (G) and M (H) cells in presence (+Q; 2 mM) or absence (-Q; 0 mM) of glutamine. Oligo, oligomycin A; FCCP, carbonyl cyanide p-trifluoro-methoxyphenyl hydrazone; Ant, antimycin A; Rot, rotenone. Data are represented as mean ± SEM of three independent cultures.

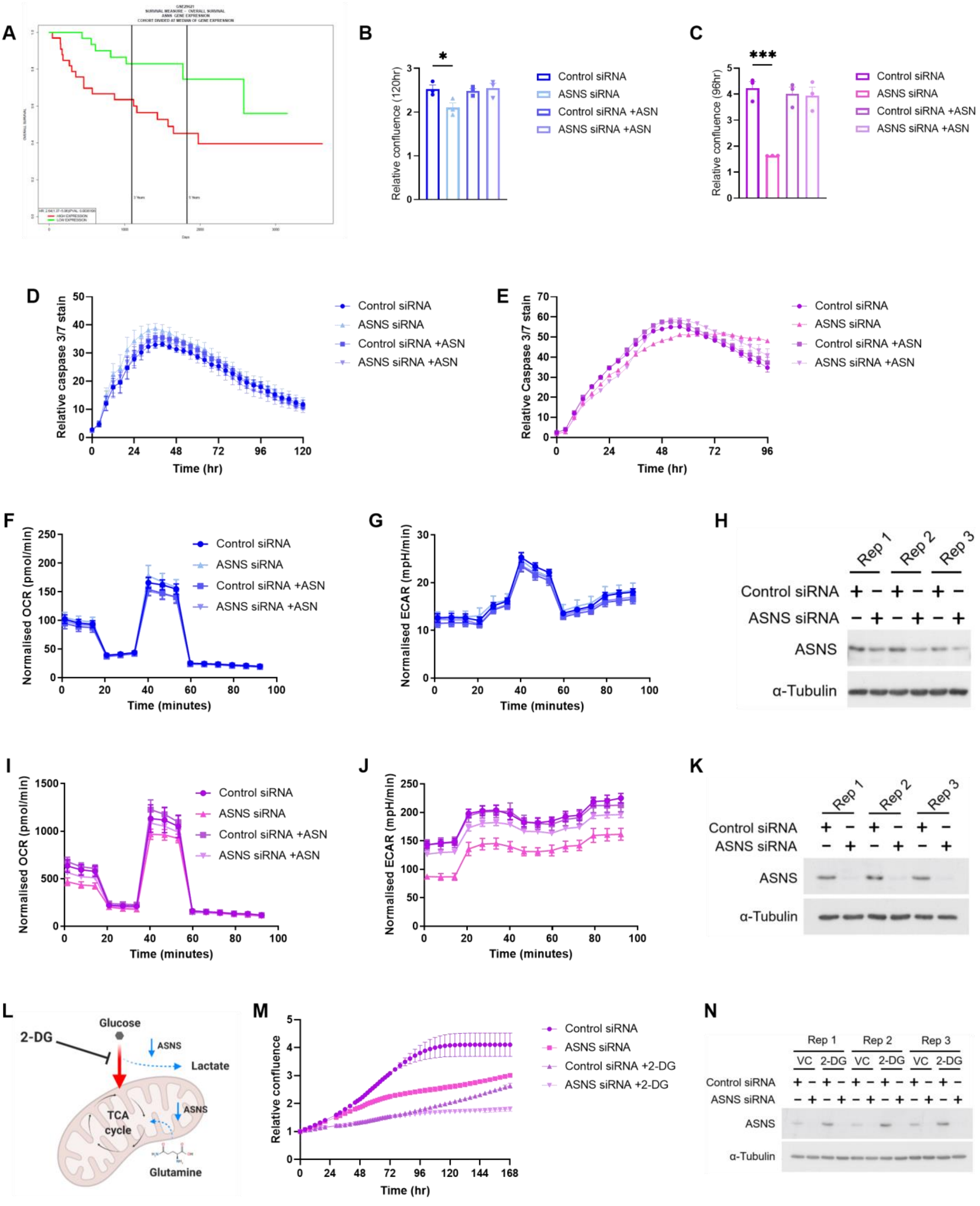


**Supplementary Figure S5. ASNS supports tumour cell phenotype**

Legend on following page

**Supplementary Figure S5. ASNS supports tumour cell phenotype**

(A) Overall survival analysis in relation to *ASNS* expression using GSE29621 and PROGgeneV2 (23). Cohort divided at median of *ASNS* expression. n=65; HR 2.64; p=0.004. (B & C) Proliferation assay in C1 (B) and M (C) cells following transfection with non-targeting control (Control) or *ASNS*-targeting siRNA, with or without 0.1 mM asparagine (ASN) supplementation. (D & E) Apoptosis assays. Cleaved caspase 3/7 staining relative to well confluence in C1 (D) and M (E) cells in assay described in (B & C). Oxygen consumption rate (OCR; F & I), extracellular acidification rate (ECAR; G & J) and immunoblots of ASNS abundance (H & K) from C1 (F-H) and M (I-K) cells following transfection with non-targeting control or *ASNS*-targeting siRNA, with or without 0.1 mM ASN supplementation. (L) Schematic of siRNA-mediated *ASNS* knockdown in M cells. *ASNS* knockdown using siRNA reduces glutamine carbon entry into the TCA cycle, causing compensatory glucose influx. This diverts glucose carbon away from lactate production, potentially allowing targeting of glucose influx using glucose analogue 2-Deoxy-D-Glucose (2-DG). Figure created using BioRender.com. (M) Proliferation of M cells following transfection with non-targeting control or *ASNS*-targeting siRNA and treatment with 10 mM 2-DG or vehicle control (VC) of water. (N) Immunoblot of ASNS abundance from experiment carried out in (M). (H, K & N) α-Tubulin serves as loading control. Rep; replicate. (A-N) Data are represented as mean ± SEM of three independent cultures. (B & C) Student's t-test. *p<0.05; ***p<0.001.

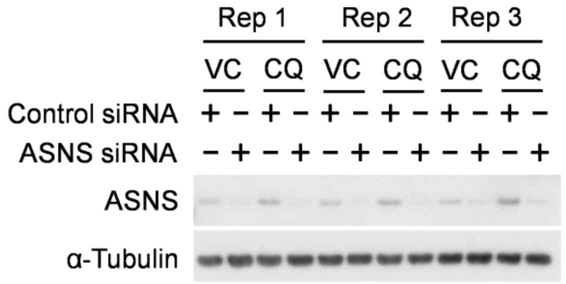


**Supplementary Figure S6. ASNS abundance in chloroquine treated cells**

ASNS abundance analysed by immunoblot following transfection with non-targeting control (Control) or *ASNS*-targeting siRNA, with or without 10 µM chloroquine (CQ). Water used as vehicle control (VC). α-Tubulin serves as loading control. Rep; replicate.
